# De Novo Genome Assembly and Characterization of Plant Growth-Promoting Rhizobacteria isolated from field grown Canola Plants

**DOI:** 10.1101/2025.04.11.645141

**Authors:** Christopher Blake, Storme de Scally, Atharva Bhide, Aysha Sezmis, Vanessa Wong, Harsh Raman, Michael J. McDonald

## Abstract

Plant growth-promoting rhizobacteria have the potential to reduce reliance on synthetic fertilizers. Yet the rhizosphere microbiome of canola (*Brassica napus*) remains understudied despite the crop’s global significance. In this study, we isolated and characterized 12 bacterial strains from canola roots, to better understand the diversity and potential agricultural benefits of the canola microbiome. Using a combination of long- and short-read whole-genome sequencing, we generated high-quality genome assemblies of all 12 bacterial species. Genomic analysis revealed genes linked to nitrogen fixation, phosphorus solubilization and phytohormone production, suggesting these bacterial strains may play beneficial roles in plant growth and resilience. Growth assays showed that most isolates proliferated in the presence of canola root exudates, indicating their adaptation to the rhizosphere. Several strains also exhibited nitrogen fixation traits, including growth in nitrogen-deficient media and ammonium production, yet bacterial inoculation did not significantly enhance early seedling development under nitrogen-limited conditions. Our findings expand the current knowledge of the diversity and functional potential of the canola microbiome and highlight promising bacterial candidates for development as biofertilizers or biocontrol agents, with implications for improving soil health and canola productivity.

## Introduction

Plants harbor a complex microbial community, termed microbiome, that influences their growth and health. The microbiome is actively shaped by the plant through the secretion of root exudates recruiting beneficial microbes, such as plant growth-promoting rhizobacteria (PGPR). PGPR enhance plant health and productivity by facilitating nutrient acquisition, producing phytohormones, and enhancing the plant’s defences against pathogens^1–4^. They have been widely studied for their ability to improve crop yields, making them valuable candidates for sustainable agriculture^5^. However, while the microbiomes of many major crops have been well-characterized, the microbial communities associated with canola (*Brassica napus*) remain relatively understudied^6,7^.

Canola is one of the most economically important oilseed crops grown worldwide. However, it has comparatively high nutrient requirements, particularly for nitrogen^8^. Traditionally, these are met through synthetic fertilizers, which has significant drawbacks, including increased greenhouse gas emissions, nitrate leaching, excess application and long-term soil degradation9–11. Plant growth-promoting rhizobacteria present a promising alternative as they can increase nitrogen availability by fixing atmospheric nitrogen into bioavailable ammonia^12,13^and producing phytohormones such as auxins, which increase the plant’s ability to absorb nitrogen from the soil^14^. Unlike legumes, canola does not form symbiotic relationships with rhizobia for atmospheric nitrogen fixation, and only a limited number of studies have reported rhizobacteria with nitrogen-fixing capabilities associated with canola^15^.

Indeed, despite the agricultural importance of canola, little is known about the genetic diversity and functional potential of bacteria inhabiting its rhizosphere.

In this study, we focus on the identification and characterization of novel bacterial isolates from the canola rhizosphere. We isolated and characterized 12 bacterial species from the rhizosphere of canola plants, representing three phyla: *Bacillota, Bacteroidota*, and *Pseudomonadota*. Using a combination of long- and short-read DNA sequencing, we generated 12 high-quality, closed genome assemblies. Further, we assessed their metabolic capabilities, including carbon utilization, antibiotic resistance and their potential to fix atmospheric nitrogen. To evaluate their potential as PGPR, we conducted growth assays in plant-associated environments, demonstrating their ability to proliferate in the presence of canola root exudates. These findings provide insights into the diversity and functional potential of plant growth-promoting rhizobacteria associated with canola, offering a foundation for developing microbial inoculants that could reduce reliance on synthetic fertilizers in canola cultivation.

## Methods Growth media

All strains were routinely grown in Tryptic Soy Broth (TSB) (15 g/L tryptone, 3 g/L soytone, 5 g/L NaCl, 2.5 g/L dipotassium phosphate, 2.5 g/L glucose) and plated on Tryptic Soy Agar (TSA): 15 g/L tryptone, 5 g/L soytone, 5 g/L NaCl, 15 g/L agar. Plants were grown in Murashige and Skoog (MS) Basal Medium (Sigma-Aldrich) with Nitrogen (M5519) and without Nitrogen (M0529) supplemented with 0.3% Agar (3g/L).

Bacterial growth was assessed using TSB, Minimal medium M9 (6.4 g/L Na□HPO□·7H□O, 1.5 g/L KH□PO□, 0.25 g/L NaCl, 0.5 g/L NH□Cl, 0.002 g/L MgSO□·7H□O, 0.011 g/L CaCl□) with either 2% glucose (20 g/L) or a 2% sugar mix (5g/L sucrose, 5g/L glucose, 5g/L mannose, and 5g/L fructose), and liquid MS with nitrogen. To detect growth on agar without nitrogen Jensen’s Medium (20 g/L sucrose, 1 g/L K□HPO□, 0.5 g/L MgSO□, 0.5 g/L NaCl, 0.1 g/L FeSO□, 0.005 g/L Na□MoO□, 2 g/L CaCO□, 15 g/L agar) was used. To measure ammonium production Burk’s Medium (0.2 g/L magnesium sulphate, 0.8 g/L dipotassium phosphate, 0.2 g/L monopotassium phosphate, 0.13 g/L calcium sulphate, 0.00145 g/L ferric chloride, 0.000253 g/L sodium molybdate, 20 g/L sucrose) was supplemented with Nessler’s reagent (Sigma-Aldrich).

### Sampling location for canola root-associated bacteria

Bacterial strains were isolated from the roots of field grown canola plants collected from two experimental field sites at Mangoplah, NSW, Australia (35.3532054°S, 147.2901734°E)) and Wagga Wagga Agricultural Institute, Wagga Wagga, NSW, Australia (35°21’09.2”S 147°17’26.5”E) on October 10, 2022. Canola plants were excavated with their root systems intact using a sterile shovel. Soil loosely attached to the roots was removed by gentle shaking, and rhizosphere soil was collected in 50 mL Falcon tubes. The taproot and main lateral roots were excised with sterile secateurs and stored in 1× Phosphate Buffer Solution (PBS).

To establish baseline soil conditions, two control samples (1A, 2A) were collected approximately 0.5 metres from plant-associated samples (3A, 3B). Soils were sampled from the 0-30 cm layer, placed in polyethylene bags prior to analysis. Soils were dried at 105°C for 24 h, lightly crushed and sieved through a 2 mm sieve. Soil pH and electrical conductivity (EC) were measured in 1:5 soil:water (w/v) extracts using a conductivity-TDS-Salinity pH-ORP-Temperature meter. Particle size distribution was determined on a Malvern Mastersizer following pretreatment with 10% H_2_O_2_ to remove organic matter. Soil organic carbon and total nitrogen were determined on finely milled samples by high temperature combustion on a Perkin Elmer 2400 Series II CHN Analyser. All samples were analysed in triplicate.

### Initial isolation and identification of bacterial strains from canola roots

Bacterial strains were isolated from lateral canola roots. The roots were placed in 25 mL of 1× PBS and vortexed for 30 seconds to remove surface soil and debris. The washed roots were then transferred to fresh 25 mL 1× PBS, vortexed for 30 seconds, and shaken for 10 seconds. This process was repeated three times.

After washing, 100 µL of the final suspension was serially diluted up to 10 □ □ and plated on TSA. Plates were incubated at room temperature for 48 hours to allow bacterial colony formation. Colonies that exhibited distinct morphologies - colour, size, and texture - were re-streaked onto TSA plates to obtain pure cultures. Single colonies were then grown overnight in 2mL TSB and stored as frozen stock with 20% Glycerol.

### Genome assembly, annotation, and assessment

Genomic DNA was extracted using the GenElute™ Bacterial Genomic DNA kit (Sigma-Aldrich) according to the manufacturer’s instructions. DNA concentration was measured using a Qubit fluorometer. Short-read sequencing was conducted using Azenta’s (Suzhou, China) generation sequencing service, which obtained paired-end 150□bp reads using an Illumina Novaseq 6000. Adaptor trimming was performed at Azenta but reads were further filtered and trimmed based on quality score using the BBDuk package (http://jgi.doe.gov/data-and-tools/bbtools/). All reads with a quality score < Q30 were removed from analysis.

In addition, long DNA fragments were selected using AMPure XP beads in a 0.4:1 ratio. DNA libraries were prepared for long-read sequencing using the Rapid Barcoding Kit 96 (SQK-RBK110.96) (Oxford Nanopore). Barcoded samples were pooled, and sequencing was performed for up to 72 hours R9.4.1 flow cells (FLO-MIN106) using a MinION sequencing device (Oxford Nanopore). Basecalling and demultiplexing were conducted using Dorado v0.1.0 (https://github.com/nanoporetech/dorado) with the super-accuracy model. Adapters and barcodes were trimmed using Porechop v0.2.4 (https://github.com/rrwick/Porechop) and reads were filtered based on quality and length using NanoFilt v2.6.0^16^.

Genomes were assembled using Flye v.2.8 ^17^, Raven v.1.1.10 (https://github.com/idaholab/raven), and Miniasm v.0.3 (https://github.com/lh3/miniasm), and the assembled genome was chosen depending on which assembler produced the most complete genome. Assemblies were polished using medaka (https://github.com/nanoporetech/medaka), Polypolish^18^, POLCA (https://github.com/alekseyzimin/masurca), and ntEdit (https://github.com/bcgsc/ntEdit). Assembly outputs were compared using dnadiff from MUMmer3 to assess genome closure and consensus accuracy^19^. If consensus sequences aligned and closed genomes were obtained, those assemblies were selected. The quality of the assembly was assessed using BUSCO v5.1.3^20^. Annotation was performed using Prokka v1.13^21^, and plant growth-promoting genes were predicted using EggNOG-mapper v2.1.12^22^. A phylogenetic tree was constructed by multiple alignments of the 16s sequence using CLUSTALW ^23^ and PhyML^24^ and plotted using ggtree^25^ in RStudio.

### Growth characteristics

Carrying capacity, area under the curve (AUC) and growth rate were determined using optical density (OD) measurements. Cultures were standardized to an equal starting CFU/mL of 10^4^ cells, based on prior CFU counts for each strain. Samples were loaded into 96-well plates in either TSB, M9 with glucose, M9 with a mix of sugars or MS with glucose in a randomized design (132 µL per well) with at least five replicates per strain. Plates were incubated at 28°C shaking, and OD was measured every 10 minutes using a plate reader (Tecan Sunrise™). Growth data were analyzed using the Growthcurver^26^ package in RStudio.

### Characterization for antibiotic resistance

Antibiotic resistance genes were identified using ResFinder^27^. To determine resistance levels experimentally, strains were grown in TSB supplemented with a gradient of antibiotic concentrations of ampicillin, chloramphenicol, ciprofloxacin, rifampicin, streptomycin and tetracycline. Samples were grown in 96-well plates in a randomized design (132 µL per well) with three replicates per strain. Plates were incubated in a plate reader (Tecan Sunrise™) maintained at 28°C, and OD at wavelength of 600 nm was measured every 10 minutes. A culture was classified as resistant if its OD increased by at least 0.07, reaching a minimum of 0.15 OD within 24 hours.

### Characterization for nitrogen fixation

Ammonium production was estimated using Nessler’s reagent. Bacterial strains were grown in 2 mL Burk’s liquid media at 28°C for 48 hours^28^. Following incubation, 80 µL of Nessler’s reagent was added to each sample, incubated for 20 minutes at room temperature, and absorbance was measured at 420 nm.

### Material Used for Growth Experiments

The seeds of Australian canola commercial cultivar, ATR-Gem bred by Nuseed Pty Ltd and Agriculture Victoria, were used for growth assays. To ensure genetic homozygousity/homogeneity, seeds were harvested from a single plant and repeatedly selfed for 6–8 generations.

### Seed sterilization

Seeds were surface-sterilized with 2% sodium hypochlorite for 6 minutes and then rinsed five times with sterile Milli-Q water before stratification in the dark for 2–3 days to promote uniform germination.

### Microbial growth in plant microcosms

To determine microbial growth in the presence of plants, microcosms were set up in square 100 mL media bottles containing 25 mL of MS medium supplemented with 0.3% agar. Three sterile canola seeds were placed on the agar and inoculated with 10 µL of either a monoculture of each species or a microbial community mixed in equal ratios at a density of 10□cells/microcosm. Microcosms were sealed with AeraSeal and maintained in a plant chamber with a 16-hour light cycle at 23°C for two weeks. To isolate microbes, plants were carefully removed from the agar with sterile tweezers and placed into Falcon tubes containing 20 mL of PBS. To remove residual agar and non-colonizing bacteria, plants were carefully washed by vortexing for 30 seconds at low speed. To recover microbes from the roots, plants were transferred to fresh Falcon tubes containing 20 mL PBS and small glass beads, then vortexed vigorously for 3 minutes. A dilution series up to 10 □ □ was prepared and plated on TSA. For mixed communities, 10 mL of the vortexed solution was centrifuged at 4,000 rpm for 10 minutes, the supernatant was removed, and the pellet was resuspended in lysis buffer (composition). DNA extraction was performed using the GenElute™ Bacterial Genomic DNA kit as previously described. PCR amplification of the 16S rRNA gene was conducted using the primers: Forward: 5’-GATCMTGGCTCAGRWTGAACS-3’; Reverse: 5’-TATTCCCYACTGCTGCCTC-3’. PCR products were purified with the Monarch Spin PCR & DNA Cleanup Kit (New England Biolabs) for subsequent sequencing and microbial community analysis. We analysed the amplicon sequencing data using a custom Python pipeline previously established in our lab^29^.

### Effect of microbes on early plant fitness

To evaluate the impact of microbial inoculants on plant growth, microcosms were established in 500 mL glass jars containing 80 mL of 0.3% MS agar without nitrogen. A single sterile canola seed was placed at the centre of each jar and inoculated with 10 µL of either a mixed bacterial culture comprised of *Klebsiella grimontii, Erwinia presicina, Enterobacter ludwigii, Chryseobacterium sp*, and *Pseudomonas rustica* in equal ratios or a monoculture of *K. oxytoca*. Two control treatments without microbial inoculants were included: one with MS medium lacking nitrogen and another with MS medium supplemented with nitrogen. Microcosms were sealed with AeraSeal and maintained in a plant chamber under a 16-hour light cycle at 23°C for four weeks. After the incubation period, plants were carefully removed with sterile tweezers, residual agar was rinsed off, and stem height, root length, and wet weight were measured.

## Results

### Genome assembly and genetic characterization of 12 bacterial isolates from the Canola rhizosphere

To expand our knowledge of the canola root microbiome, we isolated 12 different bacterial species representing three phyla: *Bacillota* (2 species), *Bacteroidota* (3 species of *Chryseobacterium*), and *Pseudomonadota* (7 species) (Fig. 1**A**). By combining long- and short-read sequencing approaches, we successfully carried out de novo genome assembly for all isolates. The genome assemblies demonstrated high quality and completeness, with an average BUSCO score of 99.6% across all isolates. Seven of the twelve genomes showed complete BUSCO scores of 100% completeness. Notably, all genomes showed very low levels of duplication (0-0.8%), except for *Bacillus sp*. which exhibited 4% duplication (Table 1).

**Table 1.**
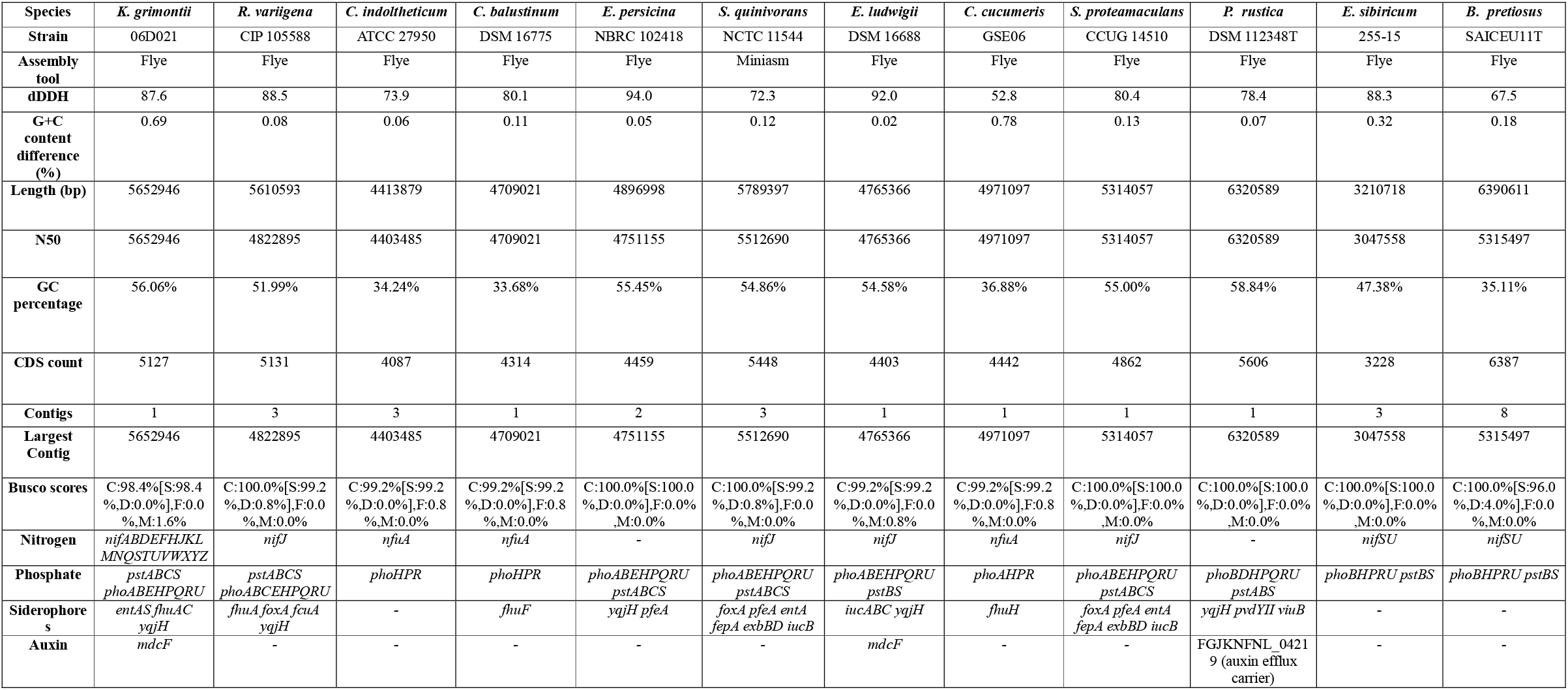
Genome assembly information.

**Figure 1:**
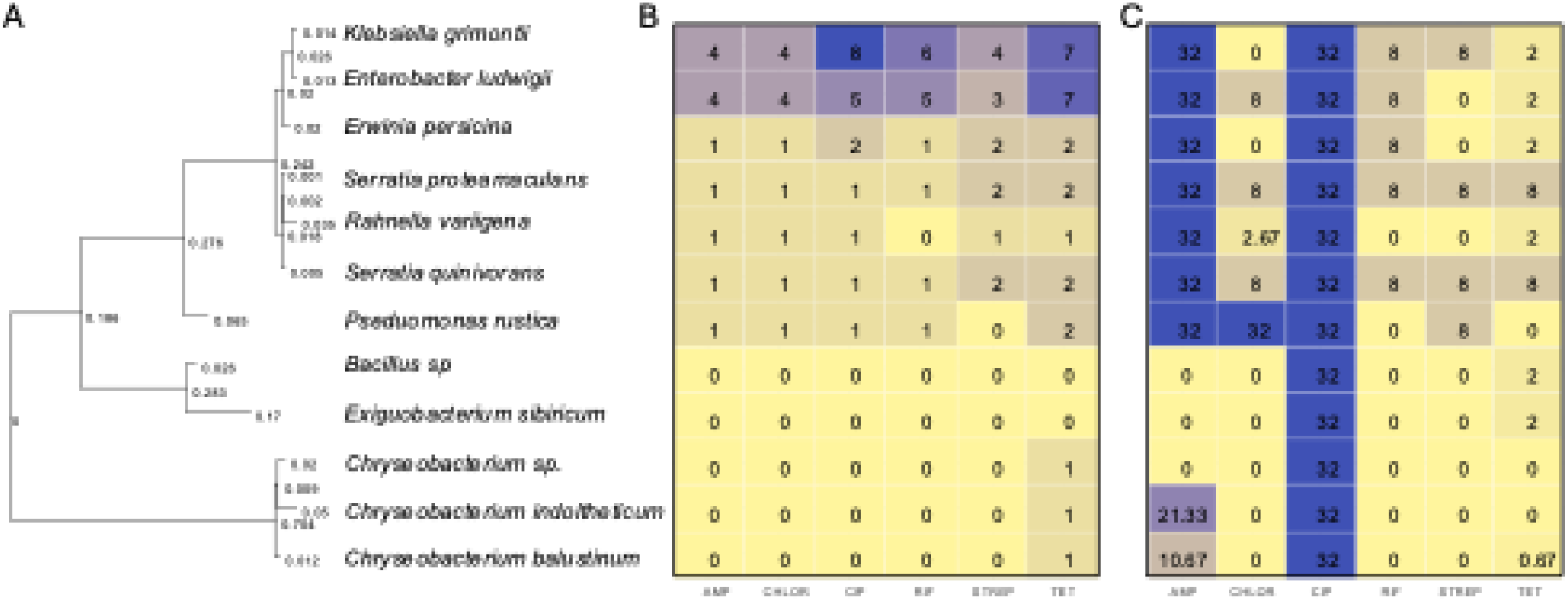
Phylogenetic tree and antibiotic resistance profiles. (A) Phylogenetic tree of bacterial strains isolated in this study, constructed based on whole-genome sequence data. The branch lengths represent evolutionary distances, with values indicating inferred genetic divergence. (B) Predicted antibiotic resistance profiles based on identified resistance genes from genome sequences. The table displays resistance scores for six antibiotics: ampicillin (AMP), chloramphenicol (CHLOR), ciprofloxacin (CIP), rifampicin (RIF), streptomycin (STREP), and tetracycline (TET). Higher values indicate a greater predicted resistance potential. (C) Experimentally determined antibiotic resistance levels from minimum inhibitory concentration (MIC) assays performed on agar plates. Values represent the MIC (in µg/mL) for each antibiotic tested. Blue shading indicates higher resistance levels, while yellow shading indicates lower resistance or susceptibility. Comment on Bootstrap values?

Genomic analysis identified four potentially novel species based on their digital DNA-DNA hybridization (dDDH) values. This computational approach serves as a replacement for traditional DNA-DNA hybridization assays and provides robust evidence for taxonomic differentiation^30,31^. The dDDH values for two species were below the 70% threshold, the established cutoff for species delineation (below which two genomes are considered distinct species): *Chryseobacterium cucumeris* (52.8%) and *Bacillus pretiosus* (67.5%). While two other isolates had values just above this threshold: *Chryseobacterium indoltheticum* (73.9%) and *Serratia quinivorans* (72.3%).

Genome sizes varied considerably among the isolates, ranging from 3.21 Mb (*Exiguobacterium sibiricum*) to 6.39 Mb (*Bacillus sp*.). Six isolates contained additional circular plasmids: *Rahnella variigena* (2 plasmids), *Chryseobacterium indoltheticum* (2 plasmids), *Erwinia persicina* (1 plasmid), *Serratia quinivorans* (2 plasmids), *Exiguobacterium sibiricum* (2 plasmids), and *Bacillus pretiosus* (7 plasmids). The N50 values closely matched the expected chromosome sizes, indicating high-quality assemblies.

The genome GC content was consistent with phylogenetic patterns, with *Chryseobacterium* species showing characteristically low GC content (33.68-36.88%), while *Pseudomonadota* members exhibited higher GC content (51.99-58.84%). The observed GC content differences between type strains and our isolates were minimal (0.02-0.78%), supporting accurate species assignments.

Genome annotation revealed varying numbers of coding sequences (CDS), ranging from 3,228 in *E. sibiricum* to 6,387 in *B. sp*., generally correlating with genome size. Our analysis detected canonical genes associated with nitrogen fixation, phosphorus solubilization, or siderophore production in some of the isolates. Interestingly, we found that many strains carried diverse antibiotic genes, which manifested as resistance to a range of antibiotics (Fig. 1**B**). Multidrug resistance can be an indication for pathogenicity in plant rhizobacteria^32,33^.

### Characteristics of field canola soil

Canola is known to modify soil properties through nutrient uptake and the release of enzymes and other root exudates^34^. The control soils (1A, 2A) were circumneutral (pH 6.47 ± 0.071) while the rhizosphere soils (3A,3B) were acidic (pH 4.31 ± 0.26) (Supplementary Data). This is consistent with previous observations of canola’s soil acidification effects, a mechanism proposed to regulate microbial activity in the rhizosphere^34^. Clay content ranged from 12.30 – 18.61 %, and the soil texture was a sandy loam^35^, which provides optimal growing conditions for canola through balanced drainage and moisture retention capabilities essential during early growth stages^36^. Soil EC was low, ranging from 137 μS/cm – 219 μS/cm, categorizing them as marginally or non-saline^37^. Carbon-to-nitrogen ratios were slightly elevated in rhizosphere soil (13.97 and 17.67) compared to control soil (9.43 and 10.06), though these values remained within previously reported ranges^38^.

### Growth characteristics of microbial isolates

To further characterize the isolates and their ability to proliferate in the rhizosphere we analysed their growth characteristics in complex and defined media. All isolates grew well in complex media at 28°C, reaching an OD> 1.0 within 24 hours (Fig. 2**A**). This indicates that the isolates can rapidly proliferate under nutrient-rich conditions.

**Figure 2.**
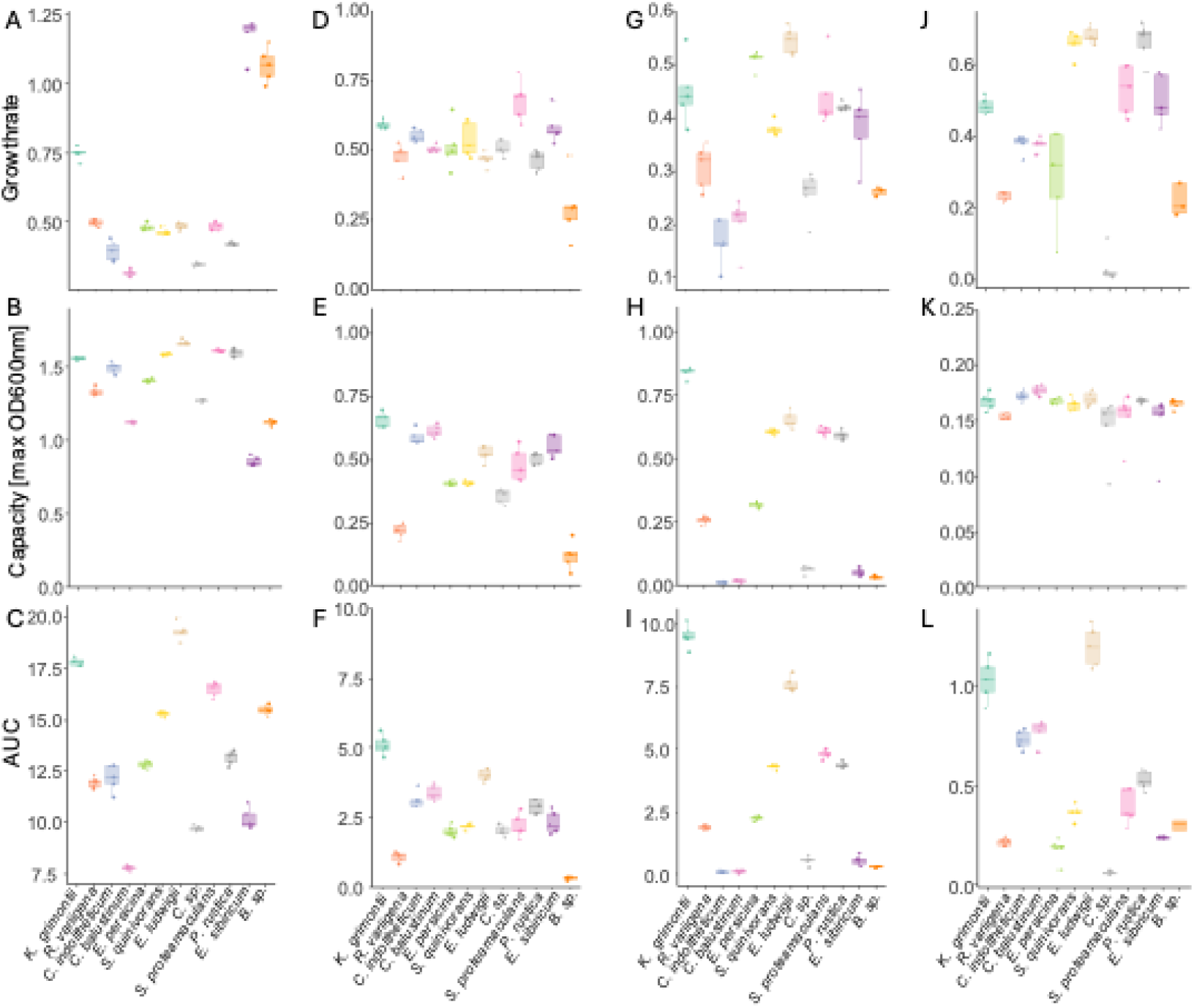
Growth characteristics of bacterial species in this study. (A–C) Growth in TSB, (D-F) Growth in M9 + 2% glucose, (G-I) growth in M9 + 2% sugar-mix, and (J-L) growth in MS +2% Glucose. Metrics include (A, D, G, J) growth rate, (B, E, H, K) carrying capacity (maximum OD600), and (C, F, I, L) area under the growth curve (AUC). Each point represents an individual replicate, and boxplots summarize variation within species.

In minimal media, all strains were capable of growth when supplemented with a single carbon source, glucose (Fig. 2**B**). Growth rates and final cell densities varied between isolates, with some exhibiting extended lag phases before entering exponential growth, suggesting differences in metabolic adaptation to glucose utilization.

To further investigate metabolic flexibility, we tested growth in minimal media supplemented with five different carbon sources (Fig. 2**C**). The growth curves revealed distinct differences, with several strains exhibiting an initial lag phase before resuming exponential growth. This pattern is consistent with a diauxic shift, where bacteria transition between preferentially utilized carbon sources^39^.

To assess whether these bacteria could grow in plant-associated environments, we tested their ability to grow in MS medium, a widely used plant tissue culture medium. None of the isolates grew in MS medium unless supplemented with an external carbon source. When glucose was provided (Fig. 2**D**), all strains showed measurable growth except *C. sp*., though at variable rates. The lack of growth in an unsupplemented MS medium suggests that these bacteria are not autotrophic and require organic carbon sources provided by the plant.

### Plant-Associated Growth in MS Medium

To assess whether bacterial isolates could grow in MS medium without an added carbon source when a plant was present, we conducted growth assays using canola seedlings (Methods). Seeds were plated on MS medium supplemented with 0.3% Agar without an exogenous carbon source, and bacterial populations were monitored over time. Colony-forming unit (CFU) counts demonstrated that most strains exhibited population growth in the presence of a canola seedling, indicating that root exudates provided sufficient organic carbon to support bacterial proliferation (Fig 3**A**). However, *C. sp, E. sibiricum* and *B. sp*. did not show detectable growth under these conditions, suggesting either an inability to utilize plant-derived carbon compounds or a requirement for additional environmental factors. We did not observe a significant impact on early seedling development and growth of either of the 12 bacterial isolates (Fig. 3**B**). Amplicon sequencing of the 16S rRNA locus confirmed that the recovered bacterial strains matched those originally inoculated. However, *S. quinvorans, C. indoltheticum*, and *C. balustinum* were below the detection limit in one or all of the replicate microcosms.

**Figure 3.**
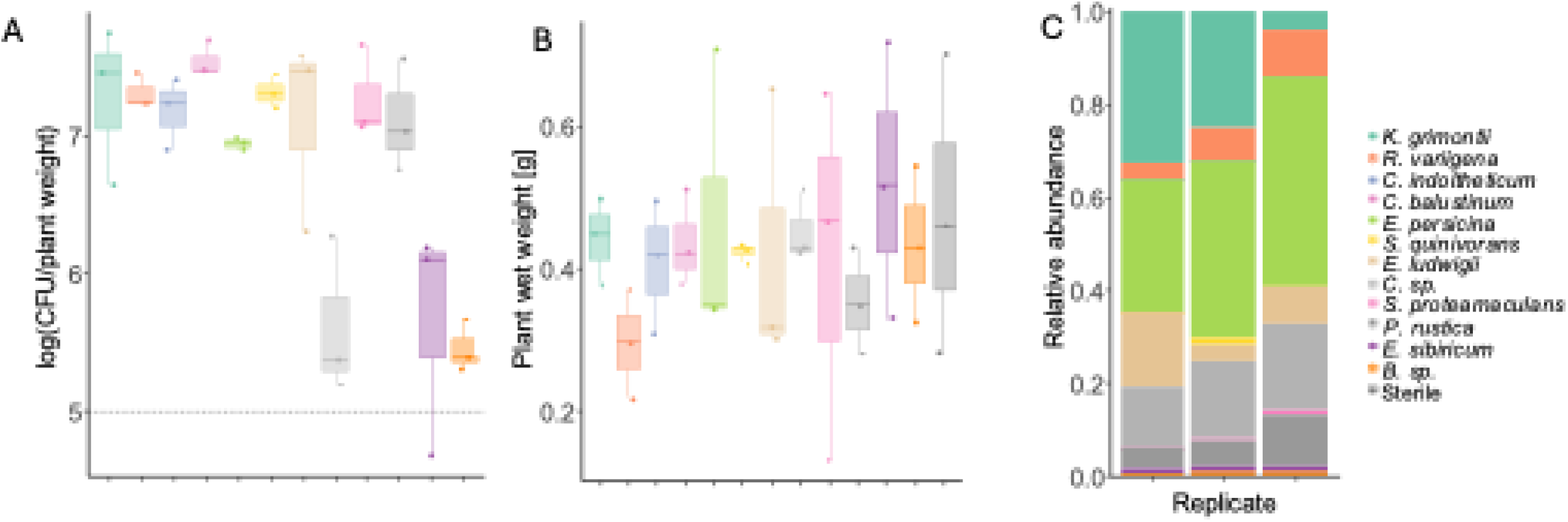
Bacterial colonization of canola seedlings after two weeks. (A) Bacterial load (log CFU per gram of plant weight) for individual bacterial strains grown in monoculture on canola seedlings. (B) Wet weight of plants after two weeks. (C) Relative abundance of bacterial species recovered from plants after two weeks. Since CFU-based measurements cannot differentiate species within a mixed community, 16S rRNA amplicon sequencing was used to identify bacterial species and their relative frequencies on three replicate plants. The composition of the bacterial community varied among replicates.

### *K. grimontii* is a nitrogen-fixing bacteria

To assess whether any of the candidate nitrogen-fixing species could fix atmospheric nitrogen, we screened the genomes of all isolates for the presence of nitrogenase-encoding genes. While several species contained genes associated with nitrogen fixation, only *K. grimontii* harbored a complete *nif* operon (Table 1). To determine whether nitrogen fixation occurred under our experimental conditions, we conducted a colorimetric assay using Nessler’s reagent, which detects ammonia production. The assay revealed that *K. grimontii* exhibited the highest OD420 values, consistent with active nitrogen fixation, while other species showed little to no detectable ammonia production (Fig. 4). These results suggest that *K. grimontii* is the primary nitrogen-fixing species in our community, potentially contributing bioavailable nitrogen to the plant-associated microbiome.

**Figure 4.**
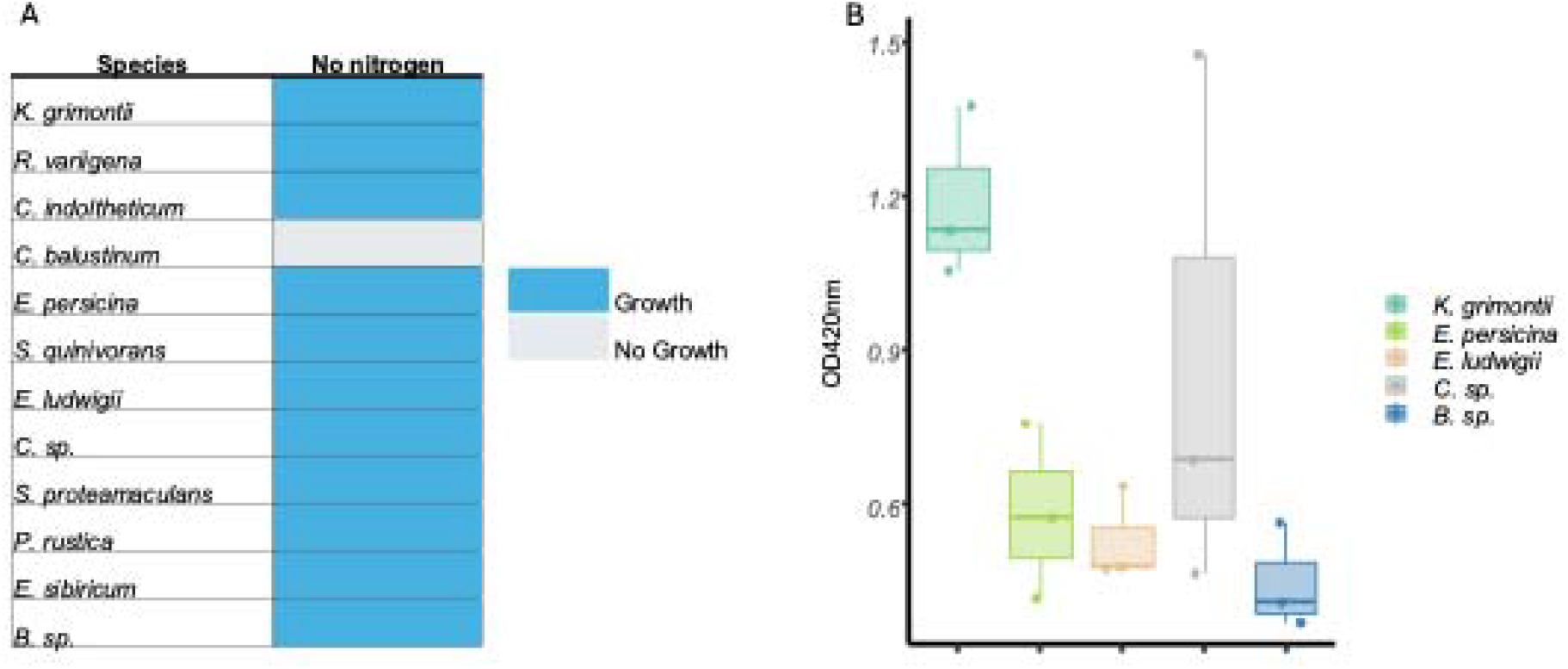
Measurement of capacity to grow without, and to fix, Nitrogen. (A) Growth of bacterial strains in M9 minimal media without nitrogen. Blue indicates growth, while grey indicates no detectable growth. (B) Ammonia production measured using Nessler’s reagent colorimetric assay. OD420nm values indicate the level of ammonia detected, with higher values corresponding to greater ammonia production. Boxplots show variation among replicates for each species tested.

### No effect of bacterial inoculation on canola seedling development under nitrogen limitation

To test if inoculation with the bacterial species can promote plant growth, we assessed the impact of bacterial inoculation on early canola seedling development under nil nitrogen conditions. Seedlings were inoculated with either *K. grimontii* in monoculture or a mixed bacterial community comprised of a subset of species (Methods) and grown without nitrogen. Two control groups without bacterial inoculation, one with and one without nitrogen, were maintained for comparison. After 4 weeks of growth, we observed a trend in increased root/shoot ratio (Fig. 5c), an indication for improved nitrogen uptake^40^, but it was not significant (p > 0.05). Notably, all plants grown in nitrogen-deficient media, regardless of inoculation status, developed purple coloration in their leaves, a characteristic symptom of nitrogen deficiency. This suggests that despite the presence of putative nitrogen-fixing genes in some *K. grimontii*, the bacterial inoculations were unable to effectively compensate for nitrogen limitation in the growth medium during early seedling development.

**Figure 5.**
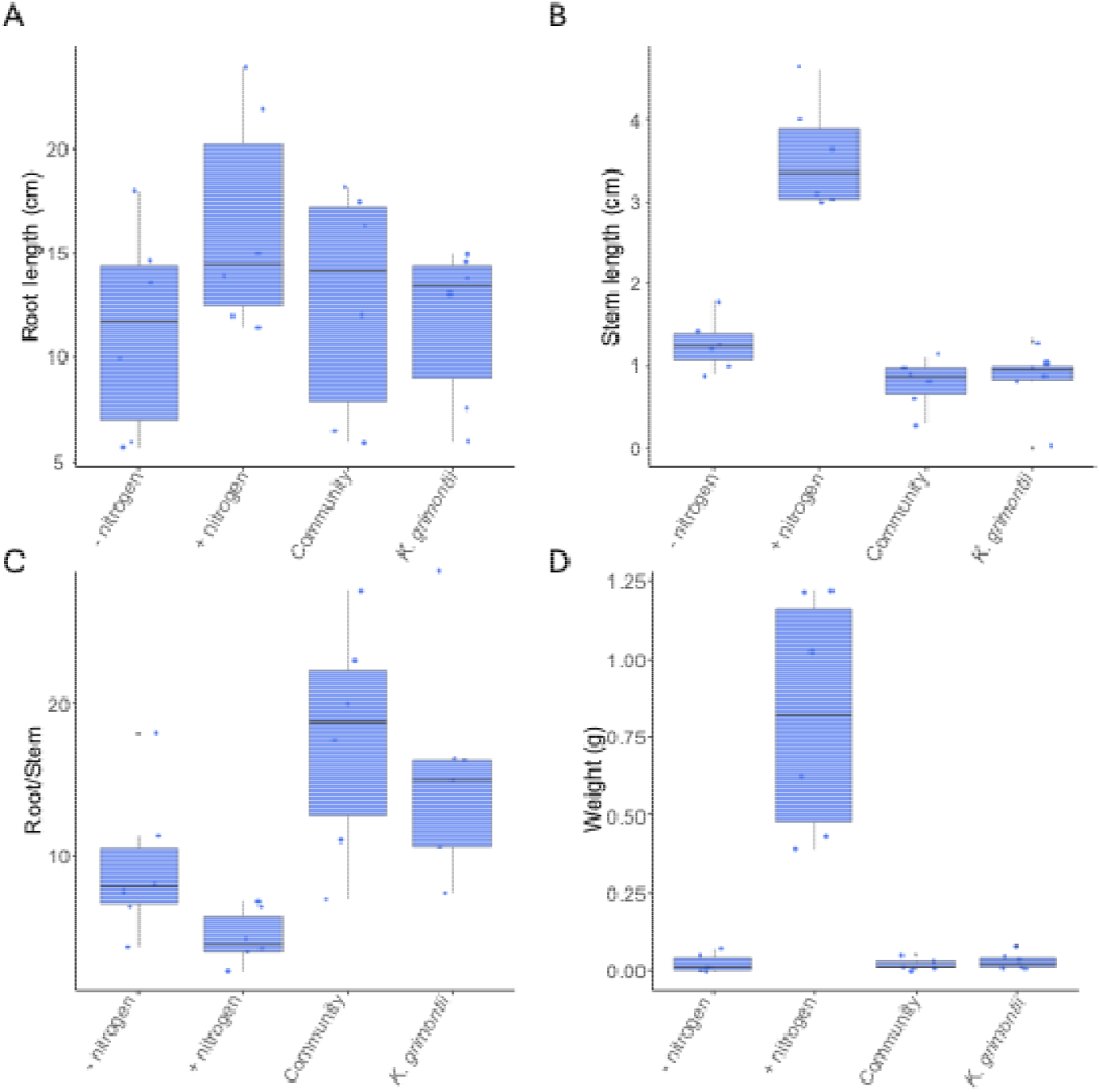
Plant growth characteristics with microbial supplementation. Measurements were taken for plants grown without nitrogen (-nitrogen), with nitrogen (+ nitrogen), without nitrogen but supplemented with a bacterial community (Community), and without nitrogen but inoculated with *K. grimontii*. (A) Root length (cm), (B) Stem length (cm), (C) Root-to-stem length ratio, and (D) Total plant weight (g). Each point represents an individual replicate, with boxplots summarizing variation within treatments. No significant differences were observed between treatments.

## Discussion

Canola is a relatively novel crop, with commercial cultivation beginning only in the 1940s. Thus, its limited history of coevolution with soil microbes suggests that we would not expect tight associations with its rhizobacteria. Previous metagenomic field studies have found either none or only one core bacterial species present in the canola rhizosphere in different seasons or sample sites^7^. Unlike well-characterized communities in model crops such as rice or maize, canola has lacked model microbial systems to investigate plant-microbe interactions at a mechanistic level. While previous research on canola-associated microbiomes has primarily relied on metagenomic approaches, our work bridges the gap between sequence-based community profiles and functional characterization^41^. By isolating and characterizing 12 bacterial strains from canola roots, we’ve established a model community that enables controlled experimental manipulation. This community now allows for systematic investigation of specific bacterial functions in the context of canola growth that could potentially reduce canola’s reliance on agricultural inputs^42^.

In modern agriculture, plant breeders have selected crop cultivars that are high-yielding and more responsive to fertilizers. While conventional agriculture addresses this challenge mainly through the application of synthetic fertilizers, this bacterial community offers the potential for the eventual development biological alternatives that could reduce environmental impacts associated with chemical fertilizers. Our genomic and experimental analyses revealed contrasting functional potential within this community: potential plant growth-promoting mechanisms alongside possible pathogenic traits. Most isolates harbour resistance genes against a wide array of antibiotic, which we confirmed experimentally. Despite antibiotic resistance sometimes indicating pathogenicity in plant rhizobacteria^32,33^ we observed no negative effect on early seedling growth and development following bacterial inoculation. Similarly, although we identified promising beneficial traits we did not detect significant improvements in canola fitness following bacterial inoculation. Canola is known for its high nitrogen requirements and low nitrogen use efficiency in commercial cultivars. Relatively few nitrogen-fixing bacteria have been reported in association with canola compared to leguminous crops^15^. In this context, our identification of a complete *nif* operon in *K. grimontii* with demonstrated ammonia production capacity is noteworthy. Several strains also contain genes involved in auxin biosynthesis pathways, which could potentially influence root development and enhance nutrient uptake efficiency. However, despite these genomic traits, we did not detect significant improvements in canola growth or root expansion following bacterial inoculation in a nitrogen-limited environment.

This disconnect between genomic potential and observable plant benefits underscores the need for deeper investigation into the factors that influence the expression and functionality of beneficial traits in realistic growing conditions. Our culturable synthetic community now provides a valuable tool for conducting such controlled experiments, allowing us to systematically explore the environmental and microbial factors governing these plant-microbe interactions. This approach will enable us to identify specific conditions that might enhance beneficial trait expression and potentially improve nitrogen fixation and plant growth in agricultural settings.

Altogether, this study adds to our understanding of canola-associated rhizobacteria and their potential as plant growth-promoting agents. By characterizing bacterial strains with beneficial traits, we provide a basis for developing microbial inoculants that could support more sustainable canola cultivation practices. Future research should focus on developing strategies to maximize their effectiveness in enhancing crop productivity.

## Data availability

Whole genome sequences generated in this study have been deposited GenBank under the Bioproject ID: <u>PRJNA1248050</u>.

## Funding

MJM was supported by ARC Discovery Grant (DP220103548).

We respectfully acknowledge Aboriginal and Torres Strait Islander peoples as the First Australians. We acknowledge the Wiradjuri people, who are the traditional custodians of the land of Wagga Wagga from where samples for this project was collected and the people of the Kulin Nations on whose land this research was conducted. We would like to acknowledge Sureshkumar Balasubramanian for introducing two of the Authors, and helping with sampling and for comments on the manuscript.

## Author contributions

CB, SS and VW carried out experiments, CB and SS carried out whole genome sequencing, CB and AS carried out genome assembly and analysis. CB carried out data visualization, made figures and all authors contributed to writing the paper.

## Competing interests

The authors declare no competing interests.

## Notes

### Competing Interest Statement

The authors have declared no competing interest.

